# Simplified artificial blood feeding and oral infection method for mosquitoes

**DOI:** 10.1101/2020.10.16.342584

**Authors:** Thiago Nunes Pereira, Fabiano Duarte Carvalho, Lidia Henrique da Silva, Silvana Faria de Mendonça, Luciano Andrade Moreira

**Affiliations:** Grupo de Pesquisa Mosquitos Vetores: Endossimbiontes e Interação Patógeno-Vetor, Instituto René Rachou - Fiocruz, Belo Horizonte, MG, Brazil

**Keywords:** Artificial blood feeding, cotton, artificial infection, Mayaro virus, *Aedes aegypti*, *Aedes albopictus*, *Culex quinquefasciatus*

## Abstract

Mosquitoes such as *Aedes aegypti, Aedes albopictus and Culex quinquefasciatus* are vectors of many pathogens that greatly affect humankind. The maintenance of these mosquitoes in laboratory permit different studies that can help understanding their biology, as well as the vector-pathogen interaction. In addition to sugar meals, the blood feeding is essential for maintenance of the reproductive cycle in several vectors. The main blood sources for many mosquito colonies are direct feeding on live animal or the use of human/animal blood through artificial feeders. However, this latter process has some disadvantages, as artificial feeders can be very laborious for assembly and decontamination. Based on these observations, a simplified technique for feeding and artificial infection was developed with cotton-pads soaked (CS) and blood or blood and viral supernatant to simulate an artificial infection. The efficiency of the CS technique was investigated through the number of mosquitoes fed/infected, when compared to their respective control group. The CS technique, with blood at room temperature, promoted a feeding rate of 61.4% for *Ae. albopictus*, 70.8% for *Cx. quinquefasciatus* and 17% for *Ae. aegypti*. The control group (Hemotec-feeding) presented 47.9%, 16.5% and 59.1% of feeding success, respectively. To improve the CS technique for *Ae. aegypti* mosquitoes, the procedure was then performed with blood at 38°C, which was possible to observe a feeding rate of 47.3%, in comparison to 53.2% for the control group (Hemotec). When using the CS technique for artificial infection with Mayaro virus, more than 80% of infection was observed for *Ae. aegypti* and 100% for *Ae. albopictus*. In the traditional infection technique (glass feeder), the infection rate was 90% (*Ae. aegypti*) and 96.6% *(Ae. albopictus*). For *Cx. quinquefasciatus*, the infection was positive only with the CS technique, resulting in 1 (5%) mosquito infected with Mayaro virus. Our results suggest that this simplified technique of low-cost feeding and easy assembly, offers good results for feeding (maintenance of colonies) and artificial infection of different species of mosquitoes.

## INTRODUCTION

Mosquitoes such as *Ae. albopictus, Cx. quinquefasciatus* and *Ae. aegypti* are vectors of several pathogens. These mosquitoes require blood feeding for obtaining proteins and egg maturation (Forattini 2002; Farnesi *et al.*, 2018, 2019). In general, the blood feeding activity of arthropods can cause problems to humans (Ajelli, 2017; Bergquist *et al.*, 2017; Benelli e Duggan, 2018), since it is during the hematophagy process that pathogens causing malaria, leishmaniasis, filariasis, dengue, zika, chikungunya, yellow fever and mayaro are transmitted (Gluber, 1998; Lehane, 2005, Cohnstaedt *et al.*, 2017; Medeiros *et al.*, 2019; Pereira *et al.*, 2020).

The maintenance of mosquito colonies under controlled conditions in laboratory is important (Pina & fonseca, 1999), allowing the development of several studies that can help understanding the biology and vector-pathogen interaction, providing foundations to direct control measures, as well as creating alternatives to manipulate the vector competence of mosquitoes (Moreira *et al.*, 2009; Landmann, 2019).

The blood feeding - beyond sugar meals - is routine to keep colonies in the laboratory, and for that, immobilized or anesthetized animals are usually used as the main blood source (Gunathilaka *et al.*, 2017; Kuo *et al.*, 2018). However, it is difficult to use animals blood source due to extensive regulations and ethical approval. Additionally, animal usage is expensive, since it needs maintenance and specialized personnel to handle the animals (Teresa *et al.*, 2011; Dias *et al.*, 2018).

Considering the importance of alternative feeding/infection methods, this study aimed to replace laborious artificial feeders (eg Hemotek and glass feeder) or direct feeding on animals. In this sense, we evaluated the effectiveness of feeding and infection of *Ae. albopictus, Cx. quinquefasciatus* and *Ae. aegypti* with an alternative technique of simplified feeding, of low cost and simplicity.

## METODOLOGY

### Mosquito species and rearing

Three species of mosquitoes were used: *Ae. albopictus, Cx. quinquefasciatus* and *Ae. aegypti*. These mosquitoes were reared in a controlled environment at 25 ± 2°C, ∼80% relative humidity and with a 12:12 light:dark. The *Ae. albopictus* and *Ae. aegypti* larvae were fed with fish food Tetramin tropical tablets^®^ and *Cx. quinquefasciatus* larvae with GoldFish color^®^. The adults, on the other hand, were fed on sugar (10% sugar solution). For experiments, the larvae were reared under the density of 100 larvae per liter of water.

### Materials and technique assembly

For the new technique, cotton was used (dental roll number 1, 100% cotton) soaked (CS) in blood at room temperature (replicates A and B) or heated to 38°C (replicate C). After being completely soaked with blood, the CS were transferred to a Petri dish and placed in the bottom of the cages (figure 1). For the infection model by the CS technique, the cotton were cut into 1 cm pieces, and placed in a Petri dish and, with the aid of a pipette, 800 μL of the viral supernatant/blood solution, maintained at 38°C, was poured on top of the cotton rods (figure 2).

**Figure 1.**
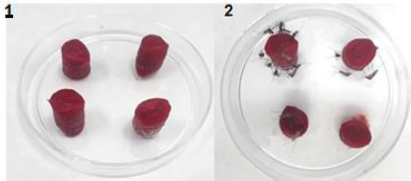
Artificial feeding of *Culex quinquefasciatus* using the CS technique (Cotton soaked). 1) The cottons were soaked in human blood, 2) separated and placed in a Petri dish 3) afterwards the cottons were placed at the bottom of the cage for mosquito feeding.

**Figure 2.**
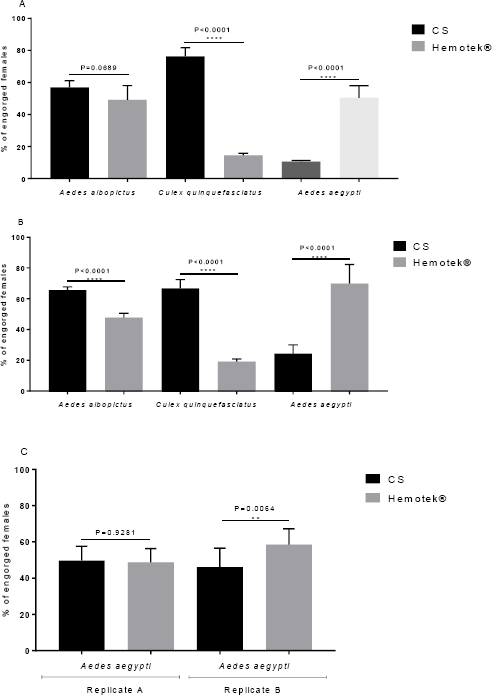
CS technique (Cotton soaked) used for artificial feeding with virus. After mixing the viral supernatant and blood (2:1), 1) cotton rods were cut into 1 cm pieces and placed in a Petri dish and moistened with 800 μL of the blood/ virus mixture. 2) the cottons were positioned at the bottom of the cage to provide the infectious feeding.

### Mosquito blood feeding experiments

Two replicates were performed at different times, and to compose the replicates, the mosquitoes were separated into 3 cages containing 100 females and 20 males for each mosquito group. Before the day of the experiment, mosquitoes were deprived of any sugar source for 24 hours. One group of mosquitoes was fed by the CS technique (blood at room temperature) and the other (control group) was fed by Hemotek^®^ which is a traditional feeder and is commonly used in many laboratories. After the results of replicates (figure 3 - A and B) two other replicas were added for *Ae. aegypti* mosquitoes (figure 3C), where the blood was heated to 38°C before offering to mosquitoes. Both treatments (CS and Hemotek^®^) were offered to mosquitoes for 1 hour and after feeding, engorged females were counted. The blood used in all experiments was obtained through a partnership between the blood bank (Fundação Hemominas) and the René Rachou Institute, under agreement (OF.GPO / CCO agreement - Nr 224/16).

**Figure 3.**
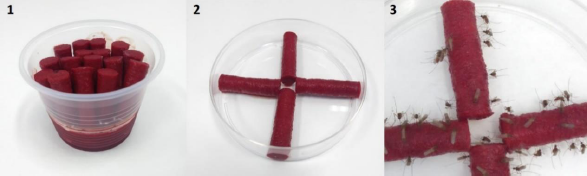
Blood feeding efficiency of the CS technique (Cotton soaked) and Hemotek^®^ feeder. Figure 3A and 3B: mosquitoes were blood fed at room temperature. Figure 3C replicate A and B: mosquitoes were fed, through the CS technique, with pre-heated blood at 38°C. The experiments were done in two biological replicates, with two independent experiments. Above each column pair we show the statistical significance, after Fischer’s exact test.

### Mosquito infection

To test CS technique for infection, we used Mayaro virus (MAYV). The MAYV vector competence has been previously tested by our group for these mosquito species (Pereira *et al.*, 2020). MAYV was grown in *Ae. albopictus* cells (C6/36), in Leibowitz L-15 culture medium, supplemented with 10% fetal bovine serum and maintained at a temperature of 28°C. The viral supernatant was collected 5 days after cell infection, quantified and frozen at -80°C until the day of the oral infection. The MAYV viral titer was 3.8 × 10^9^ PFU/mL. Adult females with 5 days of age were kept for 24 hours of fasting, and later fed for 45 minutes with a mixture of viral supernatant and blood (2:1). Simultaneously, the control groups were fed on artificial glass feeders, using a pig intestine membrane, and maintained at 38°C through a water bath. After feeding, the engorged females were separated and kept with a 10% sugar solution. All mosquitoes were collected 14 days after infection and stored at -80°C until processed.

### RNA extraction and real-time RT-PCR

Total mosquito RNA was extracted using High Pure Viral Nucleic Acid Kit (Roche), following the manufacturer’s recommendations. Thermocycling conditions were as follows: an initial reverse transcription step at 50°C for 5 min; RT inactivation/initial denaturation at 95°C for 20 sec, and 40 cycles of 95°C for 3 sec then 60°C for 30 sec. The total reaction volume was 10μL (LightCycler Multiplex RNA virus Master (Roche), 0.5μL of 10μM primers and 0.1μL of 10μM probe, and 125ng of RNA template). Each sample was run in duplicate included positive and negative controls for each virus. Viral load was determined by absolute quantification by comparison with serial dilution of the amplicon for each gene, cloned and then amplified in the pGEMT-Easy plasmid (Promega). The primers and probes used for MAYV are the according to Long *et al.*, 2011.

### Data analysis

All samples were analyzed using the Prism V 7.4 program (Graphpad). Data were analyzed using D’Agostino and Pearson’s normality test and Fisher’s exact test was used to assess differences in the prevalence of technical. The viral rate between the groups was compared using the Mann-Whitney U test.

## RESULTS

Using the CS technique (figure 3A), *Ae. albopictus* mosquitoes exhibited a feeding rate of 57% (137/240), whereas for the control group (Hemotek^®^) it was 48.7% (118/242) (P = 0.0689, Odds ratio 1,398, 95% CI, 0.976 to 2,001 - Fisher’s exact). In the second replicate (figure 3B), the CS technique promoted a feeding rate of 65.6% (166/253) while the group fed through Hemotek^®^ reached 47.7% (125/262), showing significant difference between both techniques (*P* = 0.0001, Odds ratio 2.091, 95% CI, 1.466 to 2.93 - Fisher’s exact test).

For *Cx. quinquefasciatus* mosquitoes (figure 3A), the CS technique promoted a higher successful rate [73.3% (216/283)], in comparison to the group fed on Hemotek^®^ [(14.4% of success (37/256)]. In the second replicate (figure 3B) the CS technique promoted a feeding rate of 63.5% (136/214), whereas 19.2% (39/203) in the group fed through Hemotek^®^. Both replicates exhibited statistical differences between the techniques (*P* = 0.0001, Odds ratio 19.08, 95% CI, 12.25 to 29.73 and *P* = 0.0001, Odds ratio 7.332, 95% CI, 4.691 to 11.46 respectively for replicates 3A and 3B - Fisher’s exact).

Unlike the other two species, *Ae. aegypti* showed a higher feeding rate in the control group (Hemotek^®^) in comparison to the CS technique, when mosquitoes were fed with blood at room temperature. In that case (figure 3A), the CS technique exhibited a feeding rate of 10.5% (28/265) and Hemotek^®^ showed 50.5% (134/256). In the second replicate (figure 3B), the CS technique feeding was 24.2% (57/235), while the Hemotek^®^ group presented 70% (145/207). Both replicates showed differences between the techniques (*P* = 0.0001, Odds ratio 0.1155, 95% CI, 0.07293 to 0.5636 and *P* = 0.0001, Odds ratio 0.1369, 95% CI, 0.08986 to 0.2086 respectively for replicates 3A - Fisher’s exact). Table 1 shows all replicates.

**Table 1.** Blood feeding efficiency at room temperature with two methods: CS technique (Cotton soaked) and Hemotek^®^.

| *Mosquitoes* |  | Engorged/total females |  |  |  |
| --- | --- | --- | --- | --- | --- |
|  |  | Replicate A |  | Replicate B |  |
|  |  | CS technique | Hemotek® | CS technique | Hemotek® |
| *Aedes albopictus* | A | 40/75 (53.3%) | 40/76 (52.6%) | 56/88 (63.6%) | 43/85 (50.5%) |
|  | B | 51/83 (61.4%) | 44/79 (55.6%) | 51/78 (65.3%) | 42/88 (47.7%) |
|  | C | 46/82 (56%) | 34/87 (39%) | 59/87 (67.8%) | 40/89 (44.9%) |
| *Culex quinquefasciatus* | A | 70/95 (73.6%) | 12/86 (13.9%) | 52/74 (70.2%) | 13/68 (19.1%) |
|  | B | 80/97 (82.4%) | 12/89 (13.4%) | 42/70 (70%) | 12/68 (17.6%) |
|  | C | 66/91 (72.5%) | 13/81 (16.4%) | 42/70 (60%) | 14/67 (20.8%) |
| *Aedes aegypti* | A | 9/87 (10.3%) | 36/86 (41.8%) | 14/55 (25.4%) | 39/69 (56.5%) |
|  | B | 9/90 (10%) | 46/85 (54.1%) | 16/88 (18.1%) | 47/65 (72.3%) |
|  | C | 10/88 (11.3%) | 52/94 (55.3%) | 27/92 (29.3%) | 59/73 (80.8%) |

With the low feeding rates in *Ae. aegypti* through the CS technique with blood at room temperature, two new replicates (A and B) were performed with these mosquitoes, now with blood at 38°C. With this change, the CS technique (figure 3C - replicate A), promoted a feeding rate of 49.1% (113/230), similar to the control (Hemotek^®^ - 48.5% (127/216) (*P* = 0.09281, Odds ratio 1.019, 95% CI, 0.7148 to 1,453 - Fisher’s exact). In replicate B figure 3C, the CS technique promoted a feeding rate of 45.5% (98/215), whereas the Hemotek^®^ showed 58% (137/236), statistically different results (*P* = 0.0046, Odds ratio 0.5948% CI, 04097 to 0.8637 - Fisher’s exact test).Table 2 shows all replicates with blood heated to 38°C.

**Table 2.**
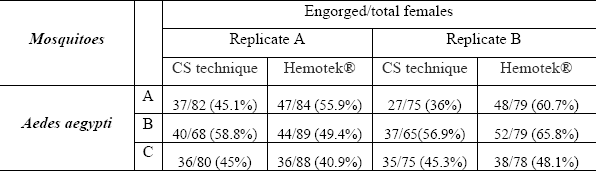
Blood feeding efficiency with blood heated to 38°C with two methods: CS technique (Cotton soaked) and Hemotek^®^.

| *Mosquitoes* |  | Engorged/total females |  |  |  |
| --- | --- | --- | --- | --- | --- |
|  |  | Replicate A |  | Replicate B |  |
|  |  | CS technique | Hemotek® | CS technique | Hemotek® |
| *Aedes aegypti* | A | 37/82 (45.1%) | 47/84 (55.9%) | 27/75 (36%) | 48/79 (60.7%) |
|  | B | 40/68 (58.8%) | 44/89 (49.4%) | 37/65(56.9%) | 52/79 (65.8%) |
|  | C | 36/80 (45%) | 36/88 (40.9%) | 35/75 (45.3%) | 38/78 (48.1%) |

When we used the CS technique to infect mosquitoes with MAYV virus (artificial infection), it was observed that at 14 days post infection, in both replicates (figure 4 A and B), *Ae. aegypti* mosquitoes exhibited an infection rate of 80% for the CS technique and 96.6% with the conventional glass feeder technique. The mosquitoes *Ae. albopictus* showed an infection rate of 100% for both techniques. On the other hand, when trying to infect *Cx. quinquefasciatus* mosquitoes, only 1 mosquito (5%) was found positive with the CS technique (figure 4A). The Mann-Whitney U test showed no difference between the groups and methods tested; on replicate A the value was *P* = 0.4543 for *Ae. aegypti, P*> 0.9999 for *Ae. albopictus* and *Cx. quinquefascitus*. For replicate B, the value was *P* = 0.7054 for *Ae. aegyti, P*> 0.8460 for *Ae. albopictus* and *P*> 0.9999 for *Cx. quinquefasciatus*. In addition, the average numbers of viral copies and infected mosquitoes remained similar between treatments (CS technique and Glass feeder), which demonstrates that CS technique does not affect the rate of mosquito infection.

**Figure 4.**
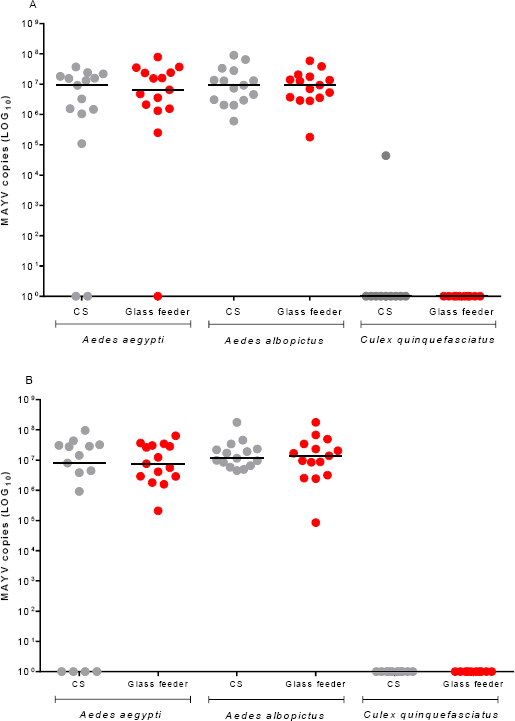
Mosquito infection with MAYV. Two infection techniques: CS technique (Cotton soaked) and Glass feeders were used to feed virus mixture to *Aedes aegypti, Aedes albopictus* and *Culex quinquefasciatus* mosquitoes, in two experiments (replicates A and B). Each circle represents a single adult female, and the black lines indicate the median number of MAYV genome copies in each treatment.

## DISCUSSION

Artificial feeding techniques for mosquitoes have been developed over the years (Costa *et al.*, 2013; Sri-in *et al.*, 2020). For example, Luo Yi-Pey (2014) calculated for *Ae. albopictus* a feeding rate of 47.9% when mosquitoes fed on pig blood using the Parafilm M^®^ method, and 42.3% when using pig blood through bovine collagen-coated membranes. The authors highlighted that these methodologies were not different for mosquitoes fed directly on mice (51.1%), which is commonly used in many laboratories for mosquito maintenance.

Ooi *et al.*, (2005) have shown that *Ae. albopictus* had a 50% success rate when fed on human blood using chicken skin and 57% when using bovine skin. Mourya *et al.*, (2000) described that *Ae. albopictus* fed with blood on a glass fiber-wrapped parafilm membrane, obtained 30 to 32% on feeding rate success. In this sense, our CS technique using human blood proved to be highly promising for *Ae. albopictus*, since we obtained a feeding rate of 61.4%.

For *Cx. quinquefasciatus*, studies have shown that, in the wild, these mosquitoes have wide food preference (Greenberg *et al.*, 2013; Hannon *et al.*, 2019), however, the feeding of *Cx. quinquefasciatus* in the laboratory usually occurs with the use of birds, making it difficult to maintain these mosquitoes. The feeding efficiency is a relevant parameter that needs to be observed in the development of new artificial feeding techniques (Costa-da-Silva *et al.*, 2013). Taking this parameter into account, our results support the conclusion that the CS technique is useful for artificially feeding *Cx. quinquefasciatus* mosquitoes, and consequently for the maintenance of these colonies in the laboratory.

Unlike the results with the previous mosquito species, *Ae. aegypti* mosquitoes presented a low feeding rate when fed with the CS technique, at room temperature. However, this low feeding rate have improved when the blood was pre-heated to 38°C before offering. Dias *et al.*, 2018 concluded that artificial feeding using citrated rabbit blood can substitute feeding on mammals, reducing the use of animals. In general, our CS technique showed satisfactory efficiency for *Ae. aegypti* mosquitoes when using heated blood. It is worth mentioning that this method does not require special materials or equipments, unlike glass feeders, which are more complex (Benedict 2009).

Regarding viral infection, our results showed that this new method proved to be effective for MAYV infection, in both *Ae. aegypti* and *Ae. albopictus* mosquitoes, with infection rates comparable to the traditional methodology. Wiggins *et al.*, 2018 have shown in an oral infection experiment, that *Ae. albopictus* mosquitoes had a significantly higher infection rate when compared to *Ae. aegypti*, a similar result observed in our study, using both methodologies. The low infectivity of *Cx. quinquefasciatus* mosquitoes may be related to their, already described, low vector competence (Brustolin *et al.*, 2018; MacLeod & Dimopouls, 2020). In addition, other studies have demonstrated this mosquito’s inability to infect and transmit Zika virus (Fernandes *et al.*, 2017; Duchemin JB *et al.*, 2017).

According to our results, we conclude that the CS technique feeding can be used to replace direct feeding on mammals or birds, reducing the use of animals, the costs of scientific research and facilitating the maintenance process of laboratory colonies, in addition to being able to be applied towards biological studies of different mosquitoes species.

## ACKNOWLEDGMENTS

We wish thankful to the Belo Horizonte Town Hall and to all members of the Mosquitos Vetores Group (MV - IRR/FIOCRUZ and World mosquito program) who helped in collecting the eggs. This work was supported by CNPq, Coordenação de Aperfeiçoamento de Pessoal de Nível Superior – Brasil (CAPES) – Finance Code 001.

## AUTHOR CONTRIBUTION

T.N.P., F.D.C., and L.A.M. contributed to the conception and design of all the experiments. T.N.P., and F.D.C., L.H.S and S.F.M were involved in the performance of the experiments. T.N.P analyzed the data and drafted the manuscript. L.A.M., F.D.C supervised the research and edited the final version of the manuscript. All authors read and approved the final manuscript.

